# Agro-morphological, yield, and genotyping-by-sequencing data of selected wheat germplasm

**DOI:** 10.1101/2020.07.18.209882

**Authors:** Madiha Islam, Abdullah, Bibi Zubaida, Nosheen Shafqat, Rabia Masood, Uzma Khan, Shahid Waseem, Mohammad Tahir Waheed, Waseem Haider, Jibran Tahir, Ibrar Ahmed, Muhammad Naeem, Habib Ahmad

**Author notes:** Corresponding authors: Ibrar Ahmed, Muhammad Naeem, Habib Ahmad.

## Abstract

Wheat (*Triticum aestivum*) is the most important staple food in Pakistan. Knowledge of its genetic diversity is critical for designing effective crop breeding programs. Here we report agro-morphological and yield data for 112 genotypes (including 7 duplicates) of wheat (*Triticum aestivum*) cultivars, advance lines, landraces and wild relatives, collected from several research institutes and breeders across Pakistan. We also report genotyping-by-sequencing (GBS) data for a selected sub-set of 52 genotypes. Sequencing was performed using Illumina HiSeq 2500 platform using the PE150 run. Data generated per sample ranged from 1.01 to 2.5 Gb; 90% of the short reads exhibited quality scores above 99.9%. TGACv1 wheat genome was used as a reference to map short reads from individual genotypes and to filter single nucleotide polymorphic loci (SNPs). On average, 364,074±54479 SNPs per genotype were recorded. The sequencing data has been submitted to the SRA database of NCBI (accession number SRP179096). The agro-morphological and yield data, along with the sequence data and SNPs will be invaluable resources for wheat breeding programs in future.

## Background and Summary

Wheat (*Triticum aestivum* L.) is the staple food crop for about 30% of the world’s population and contributes over 20% of caloric intake in diets ^1^. Current global wheat yield should be doubled to feed a projected human population of 9 Billion by 2050 ^2^. Major challenges that hamper the target of significantly increasing yield include climatic changes, reduction in arable land availability, changes in socio-economic conditions of people in developing countries, loss of biodiversity, and biotic and abiotic stresses ^3^. The target of yield increase can be achieved by investigating and utilizing the genetic diversity in available wheat germplasm, as well as improving cultivar genetics and crop management practices ^3,4^.

Genetic diversity provides a foundation for crop improvement ^5^ to develop varieties that have a better yield as well as resistance to biotic and abiotic stresses ^6^. Assessment of genetic diversity also helps to understand genomic composition, identify genes for vital traits, conserve and classify genetic variation in plant germplasm, and develop techniques for plant propagation ^6^. Since frequent use of few parents or less diverse genotypes leads to genetic erosion by producing progenies with low heterozygosity and/or inbreeding depression, it is critical to determine genetic diversity in the intended parental lines before starting a breeding program^7^. The progenies of parents with low genetic diversity may quickly become prone to biotic and abiotic stresses ^5,8^. Conversely, using diverse parental lines or genotypes can produce progenies of desirable genetic makeup, that have the tolerance to biotic and abiotic stresses, and that produce higher grain yields ^7^.

Agronomic and morphological data have been widely used to screen wheat varieties that are tolerant to stress, including drought ^9^, rust ^10–13^, salinity ^14^, and spot blotch ^15^. Molecular markers were extensively used to evaluate the genetic diversity and population structure of wheat germplasm ^16–23^. Studies using randomly amplified polymorphic DNA (RAPD) markers demonstrate narrow genetic backgrounds in most varieties introduced by the same research institutes ^16,24^. RAPD markers, however, can be problematic in terms of reproducibility and reliability, which can lead to inconsistent and/or weakly supported inferences. Single nucleotide polymorphisms (SNPs) are the most abundant polymorphism that exist in plant genomes ^25^. SNPs are appropriate for investigating marker-trait association, analyzing genetic polymorphism, mapping quantitative trait loci (QTLs), studying population structure and genomic selection. However, many SNPs are required to cover a complete genome ^26^. Recent advancements in high-throughput sequencing, not only make it possible to sequence complete organellie genomes to nuclear genomes^27–32^, coupled with the introduction of the genotyping-by-sequencing (GBS) technique has made it possible to identify genome-wide SNPs in a cost effective manner. These SNPs are useful in crop breeding, DNA fingerprinting, tagging of resistance genes for biotic and abiotic factors, and analyzing genetic diversity ^15,33–37^. For genomic DNA digestion, the restriction endonucleases utilized in GBS reduce genomic complexity, thereby enabling easier analyses of large and complex genomes such as wheat. Wheat is an allohexaploid with 42 chromosomes and has a genome size up to 17 GB ^38^. Breeders can benefit from these cost-effective informative markers during the selection of desirable wheat offspring ^39^

Among top wheat-producing countries, Pakistan ranks 4^th^ in Asia and 11^th^ in the world ^40^. To the best of our knowledge, genetic diversity in Pakistani wheat cultivars, advance lines, and landraces has not been evaluated using GBS markers. Here we report agro-morphological and yield data, along with GBS data in wheat germplasm from Pakistan. A schematic workflow of the overall study is given in Fig. 1. This data will be useful for inferring genetic diversity, population genetics, marker-assisted selection in breeding, genome-wide association studies (GWAS), mapping of rust and drought-resistant genes and other desirable quantitative trait loci (QTL) as well as for planning effective crop breeding programs in the future.

**Fig. 1.**
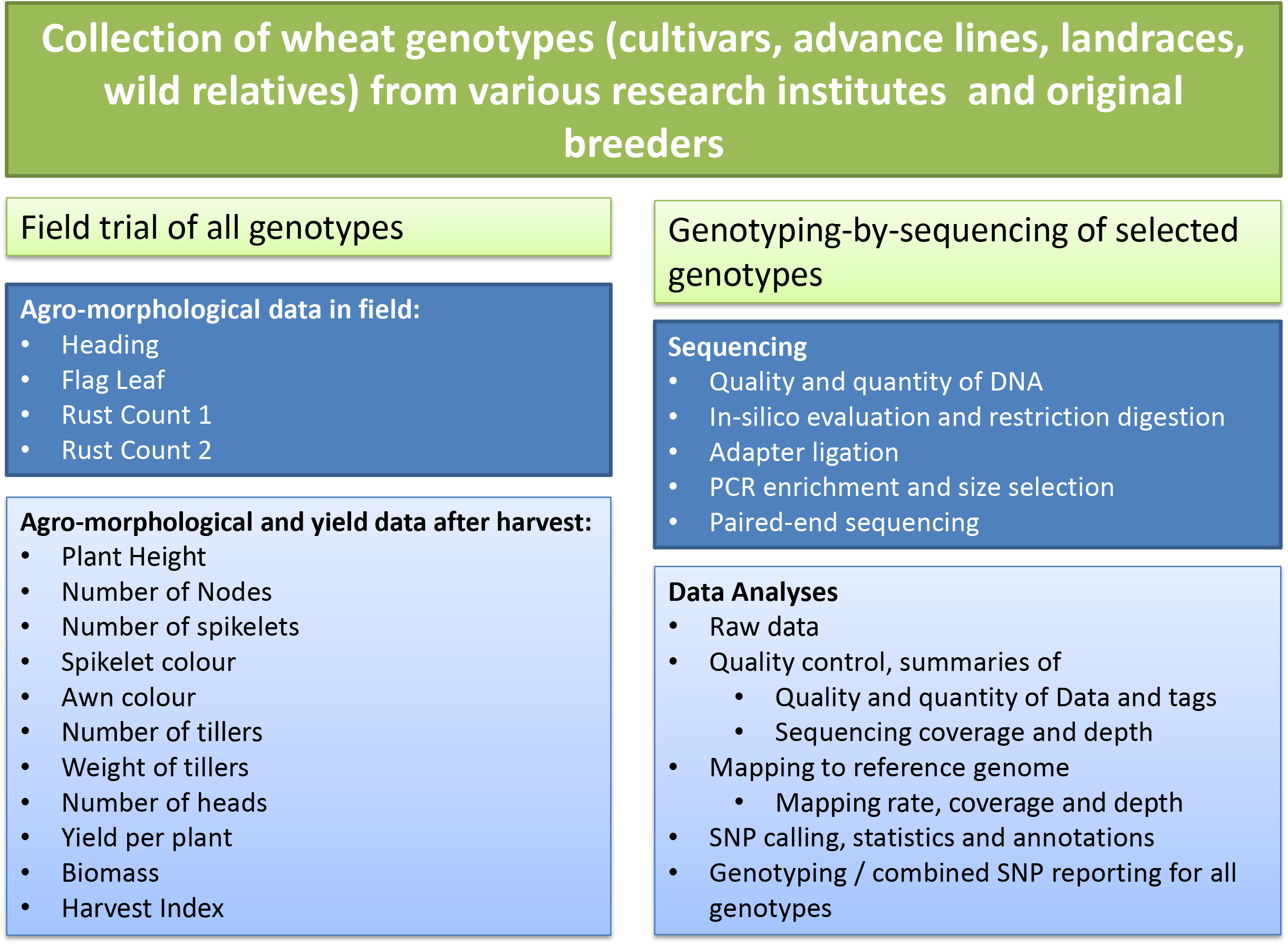
A schematic workflow of the study.

## Methods

### Collection of genotypes and field trial

A total of 104 wheat cultivars (CVs), landraces (LRs), and advance lines (ALs) were collected from different research institutes, breeders, and original collectors of landraces in Pakistan. An additional 7 cultivars were collected from separate research institutes to be included as duplicate controls in agro-morphological data. A wild relative, *Triticum monococcum* (genotype ID: 209), was obtained from the Wide Hybridization Department, National Agriculture Research Centre, Islamabad, and included in this study. Online-only Table 1 gives a list of all 112 genotypes for which agro-morphological and yield data were recorded. This table also gives the NCBI sample accession numbers of a subset of 52 genotypes, which were used to generate GBS data. Among 112 genotypes mentioned in this table, 55 cultivars are also reported in an online Wheat Atlas (http://wheatatlas.org/country/varieties/PAK/0?AspxAutoDetectCookieSupport=1; Accessed on 1^st^ August 2019). Wheat Atlas also gives details about the year of release, pedigree and selection details for these cultivars, presence of the semi-dwarf (Rht) gene, and information about the area for which the cultivar was developed. The detailed information from the Wheat Atlas for these 55 common cultivars is provided in Supplementary Table 1. The field trial was conducted in a plain field in Mandra, a town located 45 km south of Islamabad, in the Potohar region (arid zone). The geographical coordinates for the site are 33°38□N, 73°26□E. Before sowing, the field was plowed, fertilizer was homogeneously mixed in the soil, and the soil was leveled. Seeds of the genotypes were sown from 15^th^ November 2015 to 20^th^ November 2015. Each genotype was sown in one square meter block, comprising 25 plants (5 rows x 5 columns) except for four genotypes for which less than 25 seeds per genotype were available (identified in Online-only Table 1). The sixth row for all blocks comprised a rust spreader cultivar, called Morocco. The genotypes were sown in triplicate, in randomized blocks. Fig. 2 gives a snapshot of the field trial.

**Table 1.** Pearson’s Correlations among alleles of the Chakwal-50 replicates (IDs 49 and 114) using GBS data

| **Comparison among alleles** | **Pearson's Correlation** |
| --- | --- |
| Genotype 49, Allele 1, 2 | 0.997 |
| Genotype 114, Allele 1, 2 | 0.996 |
| Genotype 49, Allele 1; Genotype 114 Allele1 | 0.493 |

| **Comparison among alleles** | **Pearson’s Correlation** |
| --- | --- |
| S49, Allele 1, 2 | 0.997 |
| S114, Allele 1, 2 | 0.996 |
| S49, Allele 1, S114 Allele1 | 0.493 |

**Fig. 2.**
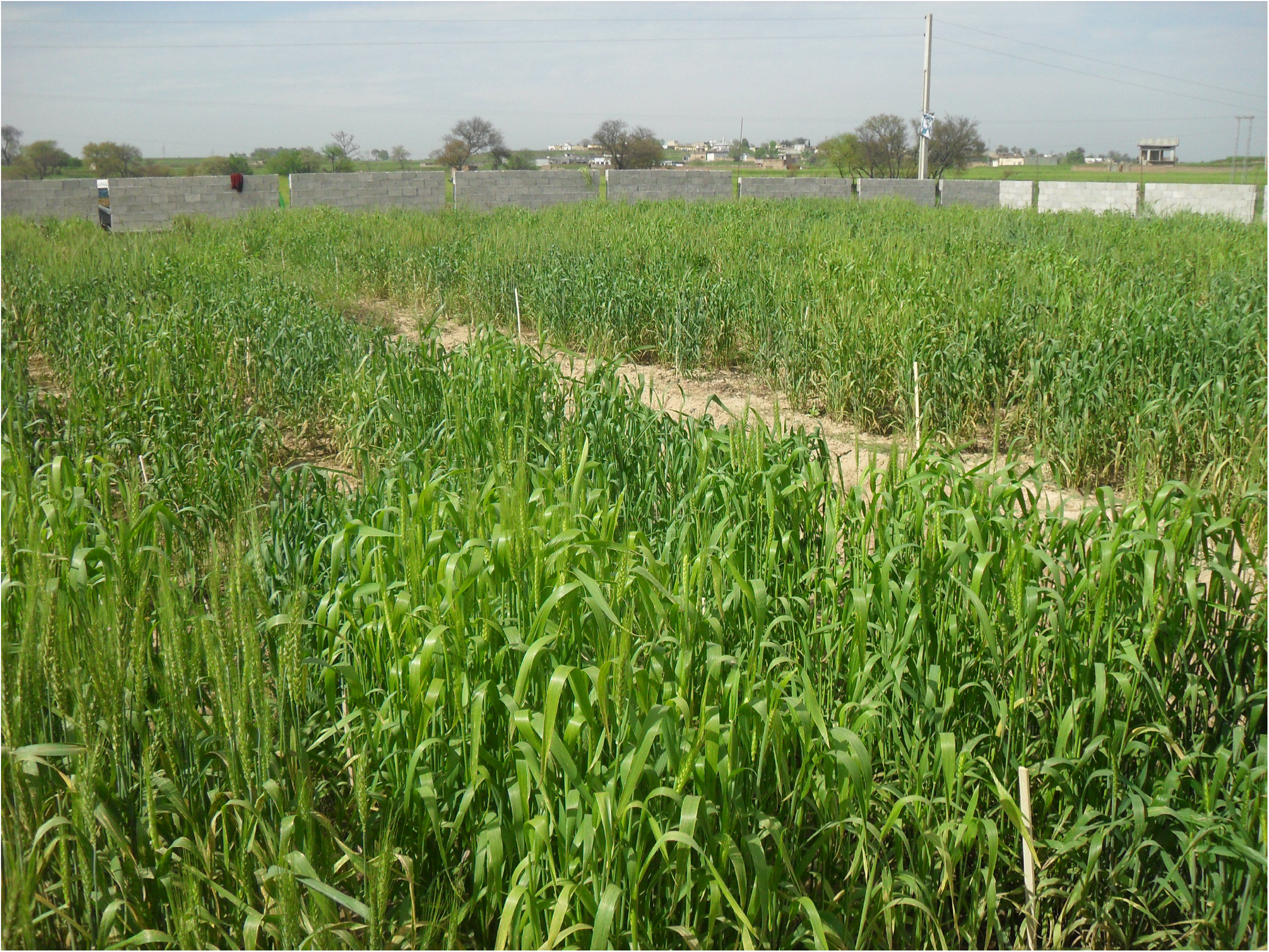
A snapshot of the field. Different genotypes visible in the field were cultivated in blocks for recording agro-morphological and yield data.

### Agro-morphological and yield data

Data were recorded in the field as well as after harvest. The field data consists of four qualitative variables. This data was based on the observation and scoring of data of entire blocks; individual plants were given the same score as that of the block for these four variables. At maturity, five plants per block were uprooted from the soil and labeled individually from 1 – 5. The plant labeling after harvest followed the EnRnPn scheme, where ‘E’ showed ‘Entry’ number (1 – 300 unique genotype IDs among 112 genotypes given in online-only Table 1), ‘R’ represented replicate number (1 – 3), and ‘P’ indicated plant number (1 – 5). For example, E1R2P5 represents entry (genotype ID) number 1, replicate number 2, and plant number 5. This labeling scheme ensured the identity of plants while recording the subsequent qualitative and quantitative data. With few exceptions, agro-morphological and yield data were recorded for 15 individual plants (five plants per replicate, in triplicates) per genotype.

### Data recorded in the field

The traits or agro-morphological variables for which qualitative data were recorded in the field included heading (H), flag leaves (FL), rust count 1 (RC1) and rust count 2 (RC2). Heading data were recorded at the booting stage for most of the plants in the field, and all data were recorded in a single field visit. The data were scored as 1 – 8, based on the presence or absence of heads on most of the plants in the entire block. Flag leaf status was recorded as drooping to erect for the entire block and given scores from 1 – 4. Stripe rust was scored on a scale from 0 - 9 as reported by Dinglasan et al ^41^. Stripe rust was scored twice; first count (RC1) was recorded 29^th^ March 2016 and the second count (RC2) was recorded on 15^th^ April 2016.

#### Data recorded after harvesting plants at maturity

After maturation, harvesting of the plants started on 30^th^ April 2016 and continued till 15^th^ May 2016. Most of the genotypes (CVs, ALs, and some LRs) were ready to harvest by the end of April; many LRs and some CVs were late in maturity and were harvested in the first and second week of May. Cold adapted LRs from the temperate region of Gilgit in northern Pakistan were the last to reach maturity. Fewer than five plants per block could be collected at maturity for these genotypes (sample IDs: 253, 255 and 256), leading to missing post-harvest data for the rest of the plants. Remaining plants for these genotypes did not reach maturity till the end of May 2016 (one month after the start of harvest) and were abandoned in the field. The following qualitative and quantitative data were recorded after the harvest:

Qualitative data were recorded for Spikelet color (SC) and Awn color (AC). The colors were scored either 1 (red to brown) or 2 (white to amber), as reported by Ormoli et al ^42^.

Quantitative data were recorded for nine variables, including Plant height (PH), Number of nodes (NN), Number of spikelets (NS), Number of tillers (NT), Weight of tillers (WT), Number of heads (NH), Yield per plant (YP), Biomass (B) and Harvest Index (HI). A brief description of each of the quantitative data recorded is given below:

1. Plant height/peduncle length (PH): Roots were cut at 2 inches from the soil. Plant height data shows peduncle length (cm) of the longest tiller from its root to the base of the spike.
2. Number of nodes per tiller (NN): Numbers of nodes were counted on the longest tiller of all individual plants.
3. Number of spikelets per spike (NS): For the spike on the longest tiller, total spikelets were counted for all genotypes.
4. Number of tillers per plant (NT): Before cutting the roots, numbers of tillers per plant originating from the same root were counted.
5. Weight of tillers (WT): After cutting the roots and spikes from all tillers, weight (in grams) was recorded. This variable represents total weight per plant excluding the weight of heads/spikes.
6. Number of heads/spikes per plant (NH): Numbers of spikes or heads were counted for all genotypes. In most cases, this number corresponded to the total number of tillers and is a measure of the number of reproductive tillers.
7. Yield per plant (YP): Seeds collectively contained in all spikes of an individual plant were threshed separately. The total weight (in grams) of the grains produced by tall spikes of one plant was recorded.
8. Biomass (B): Biomass (in grams) was calculated as the sum of the weight of tillers (WT) and yield per plant (YP).
9. Harvest Index (HI): Harvest index was calculated as the ratio of yield per plant (YP) to biomass (B), as reported by Dai et al ^43^.

### Genotyping by sequencing

Based on economic importance, a sub-set of 52 genotypes (Online-only Table 1) was selected to generate genotyping-by-sequencing (GBS) data. Seeds were grown at room temperature in plastic trays (12 inches width x 24 inches length x 2.5 inches depth; 4 x 8 cells) using autoclaved soil and sand mixed 2:1. After 14 days of sowing, leaf tissues from 10 seedlings per sample were harvested and pooled for DNA extraction using the GeneJET Plant Genomic DNA kit (Catalogue No. K0791, ThermoFisher Scientific USA). The quality and quantity of DNA were confirmed with 1% agarose gel electrophoresis and uDrop Plate of Multiskan GO (ThermoScientific, USA). DNA samples were lyophilized and shipped to Novogene Inc. Hong Kong for sequencing.

At Novogene, the purity and integrity of DNA were determined with agarose gel, and Qubit^®^ 2.0 fluorometer was used for accurate quantification of DNA concentration. For library construction, all samples contained at least 1.5 ug DNA. MseI and NlaIII restriction endonucleases were selected after *in Silico* evaluation to generate > 400,000 tags per sample and were employed for digestion of DNA (0.3-0.6 ug). Adapters were ligated to DNA along with a unique barcode for each wheat genotype. All libraries were pooled and subjected to a polymerase chain reaction (PCR) for the enrichment of sequence data. The qualified libraries were sequenced using Illumina high-throughput sequencing with 144 bp paired-end run. Average insert size of 303 bp was determined for each genotype, using Bioanalyzer.

The sequencing data was generated on a HiSeq 2500 instrument. Adapters were trimmed from the ends. Those reads which were either contaminated with library adapters, 10% unknown bases (N) or 50% low-quality bases were not used in downstream analysis. The quality of short reads was assessed using FastQC version 0.11.6 ^44^ using default parameters. *Triticum aestivum* TGACv1 ^38^ was used as a reference genome for mapping short reads using Burrows-Wheeler Alignment (BWA) version 0.7.1 ^45^ with default parameters. The reference genome was downloaded from Ensembl (ftp://ftp.ensemblgenomes.org/pub/release-33/plants/fasta/triticum_aestivum/dna; File: Triticum_aestivum.TGACv1.dna.toplevel.fa.gz; date accessed 22^nd^ March 2018).

All variants were filtered using SAMtools version 1.6 ^46^ using parameters “-q = 1, -C = 50, -m = 2, -F = 0.002, -d = 1000”. PICARD version 2.18.0 ^47^ was used to remove duplicates. To further reduce the error rate in substitutions calling, only those SNPs were selected that had coverage depth higher than 4x and mapping quality higher than 20. ANNOVAR ^48^ was used for the functional annotation of each substitution.

### Data Records

The agro-morphological and yield data are presented in Supplementary Table 2. The table also provides information about the qualitative and quantitative data for 15 plants per genotype (five plants per plot, triplicates), along with the keys used for the qualitative data. Supplementary Fig. 1 is a Box-plot representation of the dispersion in the data for all 15 variables studied. Minitab version 18 was used to generate this figure.

All GBS sequencing data and associated BAM files have been submitted in Sequence Read Archive (SRA) of the NCBI repository ^49^ and assigned SRA project number SRP179096. Individual Fastq files were given accession numbers SRR8441393 through SRR8441444; BAM files were given accession numbers SRR8467619 through SRR8467670. In total, 89.036 GB of clean data were produced; per sample data ranged from 1.01 to 2.5 GB. The lowest Phred score value for Q30 was 89.41%. The values of GC content in individual samples ranged from 42.14% to 44.17%. Information about individual samples, quantity, and quality of generated data are provided in Supplementary Table 3 along with details of each wheat variety, numbers of bases generated per sample and their respective quality values. Reference genome mapping information is given in Supplementary Table 4. This table provides a summary statistic of the mapping of short reads to the wheat reference genome.

Online-only Table 2 gives statistics about the variants called (SNPs) for individual genotypes. This table also gives functional attributes of the SNPs and gives the number of transition and transversion mutations. The average number of SNPs per genotype was 364,074 ± 54,479. When SNPs for all genotypes were merged, the total number of SNPs reached 2 Million. These combined SNPs, with exact nucleotide positions on the wheat reference genome, are given in the file “Genotyping and SNPs data” ^50^, available on Figshare. This file contains a complete record of SNPs. The data in each column can be read from left to right - #Chromosome: Chromosome position along the small arm and long arm of the chromosome, #Position: The coordinate position of nucleotide base which showed substitution, #Reference: The nucleotide present in the reference genome, #Allele: The type of substitution in the reference genome showing first the allele present in the reference genome and then the allele present in the sample sequence in the current study, #Gene: The name of the gene in which the substitution exists, #Annopos: Type of substitution according to the location, such as intergenic, genic, intronic, UTR, synonyms and non-synonyms. The next column shows the substitution present in each sample in a diploid form such that GG represents the homozygous condition and AG represents the heterozygous condition.

Studies of genetic diversity, population genetics, phylogenetics ^51–54^, association mapping and genome-wide association studies ^15,55–58^, linkage map and quantitative trait loci (QTL) mapping, marker-assisted and genomic selection ^35,63^ have used GBS data for the advancement of breeding in various plant species including wheat. Together with agro-morphological and yield data, GBS data generated for wheat genotypes in this study will be extremely useful in future crop breeding programs. The data will be helpful in the breeding of elite wheat cultivars having high yield and resistance to biotic and abiotic stresses to feed the growing human population.

### Technical validation

Seven cultivars were included as duplicate controls in the current study. These include Sahar (Genotype IDs: 37 and 143), Faisalabad 2008 (Genotype IDs: 38 and 140), Lasani 2008 (Genotype IDs: 39 and 144), Marvi 2000 (Genotype IDs: 46 and 147), Chakwal 50 (Genotype IDs: 49 and 114), Galaxy (Genotype IDs: 54 and 141), and TD-1 (Genotype IDs: 52, 131). One replicate for these genotypes (genotype IDs: 37, 38, 39, 46, 49, 52, and 54) was collected from Cereal Crops Research Institute (CCRI), Pirsabak, Nowshera, while the second replicate was collected from different research institutes: genotype 114 from Barani Agricultural Research Institute (BARI), Chakwal; genotypes 140, 141, 143, and 144 from Federal Seed Certification and Registration Department (FSC&RD), Khanewal and genotypes 131 and 147 from Nuclear Institute of Agriculture (NIA), Tandojam. These duplicated genotypes were randomly assigned separate genotype IDs and sown in the field like other genotypes. Their agro-morphological and yield data were subjected to multivariate analyses including principal component analysis (PCA) and hierarchical cluster analysis or dendrogram (Fig. 3) in Minitab version 18. Fig. 3a is the PCA plot using average values of 15 samples per genotype, and Fig. 3b shows the dendrogram of these seven genotypes. These figures show that, except for Chakwal 50 (Genotype IDs: 49 and 114), all duplicated genotypes tend to cluster together, but appear distinct from other cultivars. This observation attests to the authenticity of the agro-morphological data. Chakwal 50 replicates (Genotype IDs: 49 and 114) appeared very distinct from each other (Fig. 3a and Fig. 3b). They tend to cluster with other genotypes rather than clustering together. Due to discordance in morphological results among the replicates, both replicates (49 and 114) were subsequently selected to generate GBS data.

**Fig. 3.**
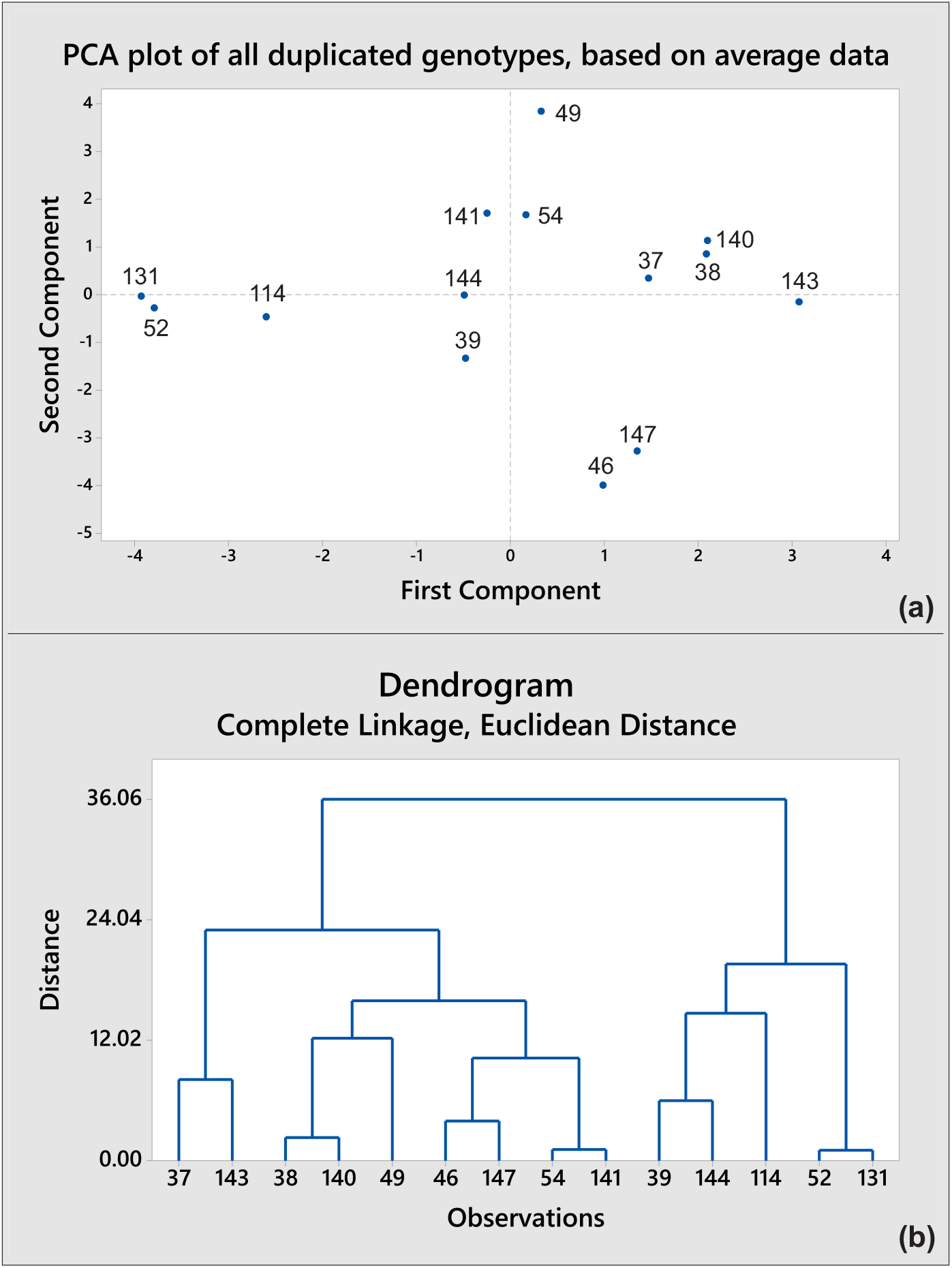
Results of the multivariate analyses, showing clustering of the duplicated genotypes. Average data of all plants per genotype was used for these analyses. The duplicate genotypes (IDs in brackets) include Sahar (37 & 143), Faisalabad 2008 (38 & 140), Lasani 2008 (39 & 144), Marvi 2000 (46 & 147), Chakwal 50 (49 & 114), Galaxy (54 & 141), and TD-1 (52 & 131). Except for the genotype Chakwal-50, both the PCA plot (a) and dendrogram (b) tend to cluster together the duplicates in each genotype.

Using the SNPs generated from the GBS data (provided in Genotyping and SNPs data file ^50^), values of Pearson Correlations between the two alleles within each replicate and between the two replicates were calculated using Minitab version 18 (Table 1). Almost perfect correlations between alleles 1 and 2 in each genotype (above 0.99 correlation values) reflected high genomic homozygosity within each replicate, nullifying the chances of mixing distinct genotypes in original DNA extractions intended for GBS. On the other hand, moderate correlations (less than 0.5) between the two replicates revealed distinctness between them. Together with the agro-morphological findings, the two replicates used for Chakwal-50 were two distinct genotypes rather than the two replicates of a single genotype. The exact identification of these two genotypes (49 and 114) could not be established from current data.

For generating GBS data, DNA from the selected genotypes was extracted after mixing young fresh growing leaves of 10 seedlings per genotype, to ensure that the sequencing data was representative of the genotype and not any individual plant. High quality of the sequencing data was evident in FastQC analyses; up to 90% of all the short reads exhibited a Phred quality score of Q30 (99.9% correct base calling) or above. SNPs were called for variants having a minimum of tag4 value (coverage depth of 4 or more) and a quality score of Q20 (99% accuracy) or more. Thus, only high-quality variants were included in the dataset. The authenticity of GBS data was evaluated by selecting 16,000 SNPs from the Genotyping and SNPs data file ^50^. These SNPs were selected using the following criteria: (a) presence of alleles in all genotypes (zero missing data), (b) common alleles among the genotypes (more than 0.3 minor allele frequency), and (c) representation of SNPs from all 42 chromosomes in the wheat genome. From these SNPs, a dendrogram was generated in the R program to show the relationship among the genotypes. Two distinct clusters were evident in the dendrogram whereby genotypes belonging to different sources of collections tended to cluster together (Fig. 4). This approach not only validated the usefulness of GBS data but also made obvious the genetic distinctness of the genotypes collected by the institutes (original sources of sample collection for this study).

**Fig. 4.**
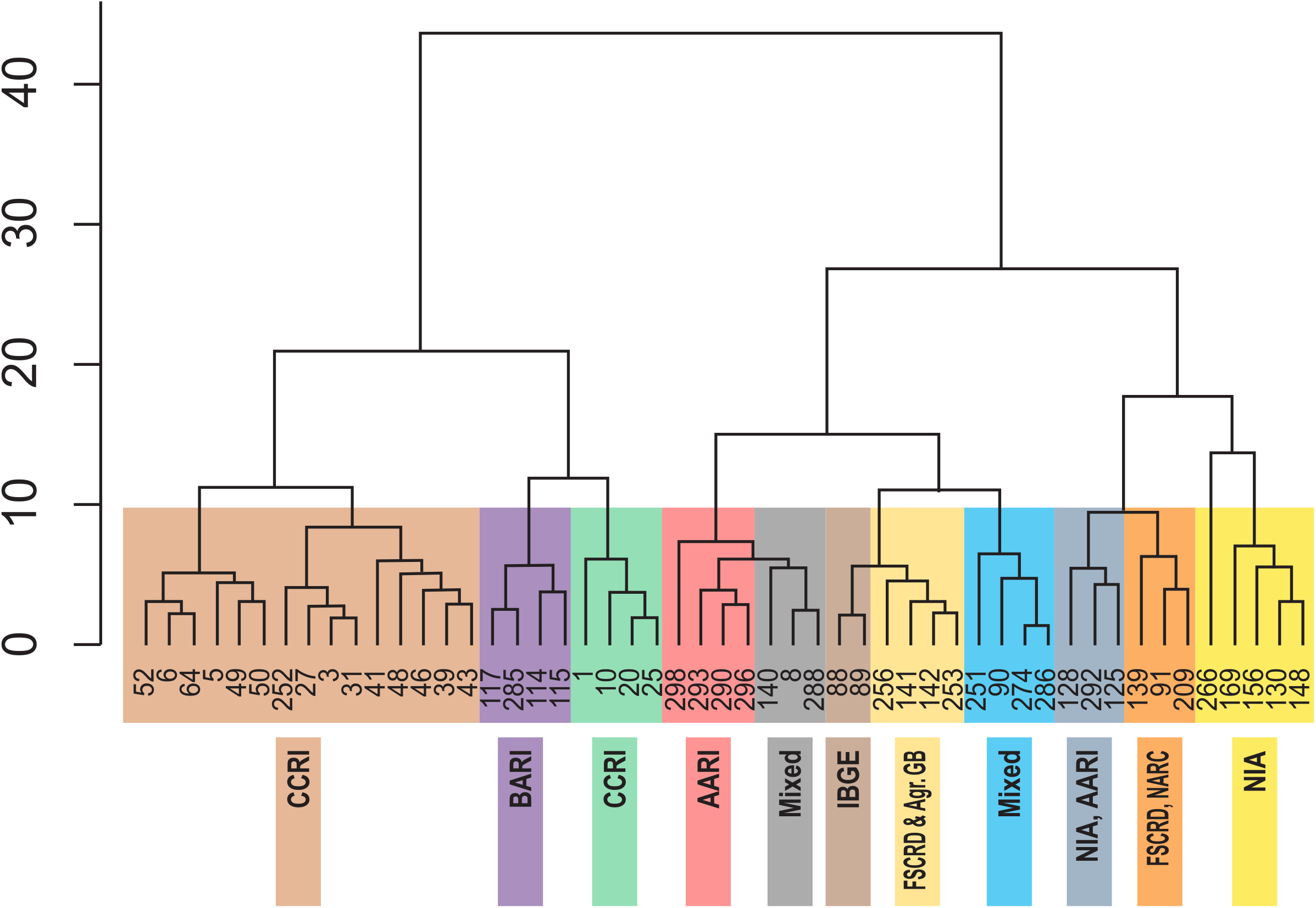
Dendrogram based on Ward distances, grouping the genotypes into clusters and subclusters. Genotypes collected from individual research institutes tend to cluster together.

### Availability of the wheat genotypes

Sources of the wheat genotypes collection have been listed in Online-only Table 1. The source institutes are expected to annually refresh and retain the propagating material, which is essential for its viability over the years. As per the Plant Breeders’ Rights Act 2016 in Pakistan, original breeders of the cultivars and advance lines retain the property rights of their breeding material. In line with this Act, the authors are not authorized to share and disseminate the genotypes covered by the Act. The authors welcome queries from other researchers and potential breeders about the availability and sharing of the genotypes which are not protected by the Act. Where applicable, the respective laws of donor and recipient countries will govern the transfer of the propagating / living material to other countries outside Pakistan.

### Code availability

All software tools used to analyze the NGS data are free to use and publicly available.

## Supporting information

Online Only Table 1

Online Only Table 2

## Acknowledgments

The authors thank contributors to the source materials used in this study. The authors thank the support and help provided by the field staff during field trials. The authors are thankful to Claudia Henriquez from Missouri Botanical Gardens, USA, for proofreading this manuscript, and the anonymous reviewers for their invaluable suggestions in improving the manuscript. The first author is a recipient of the Indigenous Ph.D. Fellowship by the Higher Education Commission of Pakistan.

## Author contributions

H.A., M.N., and I.A designed and jointly supervised the study. M.I. and M.N. collected genotype seeds from the research institutes. M.I., Abdullah, S.W., M.T.W., I.A., and M.N. conducted field trials and recorded field data. M.I., B.Z., N.S., R.M., U.K., I.A. and M.N. recorded the agro-morphological and yield data after the harvest. M.I., Abdullah, I.A., and M.N. prepared samples for GBS and extracted DNA. M.I., J.T., W.H., and I.A. analyzed GBS and genotypic data. Abdullah and I.A. submitted GBS data on NCBI. M.I. and Abdullah drafted the manuscript with input from H.A., I.A., and M.N. All authors read and approved the final version of the manuscript.

---

**Supplementary Fig. 1.** Supplementary Fig. 1 is a Box-plot representation of the dispersion in the data for all 15 variables studied.

