## Supplementary figures and images for "Agro-morphological, yield, and genotyping-by-sequencing data of selected wheat germplasm"

### Supplementary Figure 1

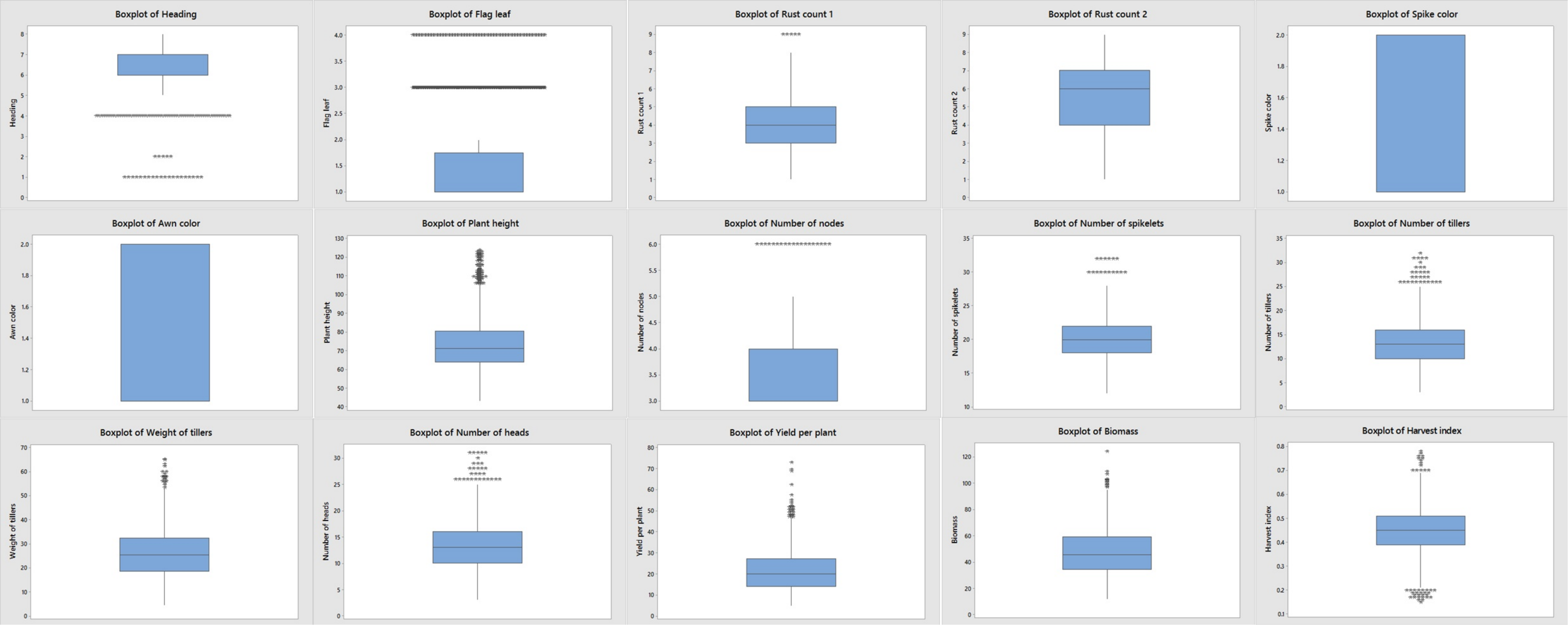
